# Key genes and pathways in asparagine metabolism in Alzheimer’s Disease: a bioinformatics approach

**DOI:** 10.1101/2025.04.25.650586

**Authors:** Xiaoqian Lan, Guangli Feng, Qing Li, Yuting Shi, Shiyi Qin, Lianmei Zhong

## Abstract

**Background:** Asparagine (Asn) metabolism is essential for maintaining cellular homeostasis and supporting neuronal energy demands. Recent studies have suggested its dysregulation may contribute to Alzheimer’s disease (AD) pathogenesis; however, the specific genes and regulatory mechanisms involved remain incompletely understood.

**Methods:** Four publicly available microarray datasets (GSE5281, GSE29378, GSE36980, and GSE138260) were utilized to investigate genes with differential expression between control and AD samples. Asparagine metabolism-related genes (AMGs) were retrieved from the GeneCards database, and their intersection with DEGs yielded candidate asparagine metabolism-related differentially expressed genes (AMG-DEGs). Functional enrichment analysis (Gene Set Enrichment Analysis, Gene Ontology and Kyoto Encyclopedia of Genes and Genomes), protein–protein interaction (PPI) network analysis, and centrality scoring identified hub genes. Regulatory mechanisms were investigated through construction of competing endogenous RNA and transcription factor networks. Potential therapeutic compounds were predicted via drug–gene enrichment and evaluated using molecular docking simulations.

**Results:** Thirty-nine AMG-DEGs were identified and found to be enriched in neurodevelopmental, synaptic transmission, and inflammatory signaling pathways. PPI analysis and centrality screening revealed seven hub genes (*HPRT1*, *GAD2*, *TUBB3*, *GFAP*, *CD44*, *CCL2*, and *NFKBIA*). Regulatory network analysis highlighted specific miRNAs, long non-coding RNAs, and transcription factors involved in their modulation. Drug screening and docking identified Bathocuproine disulfonate, DL-Mevalonic acid, and Phenethyl isothiocyanate as promising compounds with strong binding affinities to hub proteins.

**Conclusion:** This study comprehensively maps the dysregulation of asparagine metabolism in Alzheimer’s disease and reveals a set of hub genes and regulatory elements potentially involved in disease progression. The predicted therapeutic compounds provide a foundation for further experimental validation and may contribute to the development of novel metabolism-targeted strategies for AD treatment.

## 1. Introduction

Alzheimer’s disease (AD) is a chronic neurodegenerative condition marked by progressive cognitive impairment, memory deterioration, behavioral abnormalities, and eventual loss of autonomy(1, 2). Pathologically, AD is characterized by the deposition of extracellular β-amyloid (Aβ) plaques, the formation of intracellular neurofibrillary tangles consisting of hyperphosphorylated tau, and extensive synaptic dysfunction(3). Due to increasing global longevity, AD has become a leading cause of morbidity among older adults, with over 50 million cases reported worldwide—a number projected to triple by 2050(4, 5). This growing burden underscores the imperative for novel therapeutic strategies and effective preventive measures.

Emerging evidence suggests that metabolic disturbances are key contributors to AD pathogenesis beyond classical amyloid and tau pathology(6). Patients with AD often exhibit systemic dysregulation of glucose utilization, lipid processing, and amino acid metabolism(7), which may trigger neuroinflammation, synaptic breakdown, and vascular impairment(8). These metabolic disruptions can modulate transcriptional programs through regulators such as NF-κB and Nrf2(9), and may also alter non-coding RNA networks, further exacerbating neuronal dysfunction and disease progression(10). Among multiple metabolic abnormalities, altered amino acid metabolism—especially involving asparagine (Asn)—has drawn increasing research attention. Asn, synthesized by asparagine synthetase and catabolized by asparaginase, is implicated in cellular immunity(11) and diverse neuronal functions, including bioenergetics, neurotransmitter regulation, and protein post-translational modification(12). Observations from both sporadic and autosomal-dominant AD brain tissues demonstrate disrupted Asn homeostasis, suggesting a common metabolic signature associated with disease progression(13). Clinical studies have shown that plasma Asn concentrations are markedly decreased in AD patients and positively correlate with neurodegeneration-associated markers such as neurofilament light chain, particularly among individuals with elevated cerebral Aβ load(14). Mechanistically, Asn can be converted to aspartate via transamination, thereby entering the tricarboxylic acid (TCA) cycle and contributing to cellular energy metabolism(15). Metabolomics studies have corroborated these findings, showing significantly decreased Asn levels in the plasma of AD patients, which may contribute to an energy-deficient and neurotoxic microenvironment(16). Asn is also essential for protein N-glycosylation, a key post-translational modification process. Depletion of Asn may disrupt glycosylation events(17); notably, aberrant glycosylation at the Asn368 site of tau protein has been associated with impaired synaptic signaling and reduced plasticity, potentially intensifying AD-related neuropathology(18). Although asparagine endopeptidase (AEP) does not participate directly in Asn metabolism, many of its substrate proteins include Asn residues, suggesting its activity may be modulated by Asn availability(19). AEP has been implicated in the proteolytic cleavage of both amyloid precursor protein and tau, leading to the generation of neurotoxic fragments(20, 21) and influences microglial activation(21), and inflammatory cascades that impair neuronal integrity and cognitive performance(22). Although existing studies support a link between altered Asn metabolism and AD, the molecular mechanisms underlying this association remain poorly understood. In particular, the regulatory interactions of asparagine metabolism-related genes (AMGs)—especially those involved in neuroinflammation and synaptic integrity—and their potential as therapeutic targets require in-depth, systematic investigation.

This study employed bioinformatics approaches to systematically analyze the characteristics of Asn metabolic dysregulation in AD. By integrating multiple public databases, we successfully identified asparagine metabolism-related differentially expressed genes (AMG-DEGs). Furthermore, we elucidated how these genes participate in the molecular mechanisms underlying AD pathogenesis by regulating key biological processes, including neuroinflammation, synaptic dysfunction, and energy metabolism. Through functional enrichment analyses, such as Gene Set Enrichment Analysis (GSEA), Gene Ontology (GO) and Kyoto Encyclopedia of Genes and Genomes (KEGG), we systematically characterized the biological functions of Asn metabolism-related genes. Additionally, by constructing protein-protein interaction (PPI) and gene regulatory networks, we identified several hub genes that play critical roles throughout the progression of AD. Notably, through drug screening and molecular docking simulations, we identified multiple high-affinity small molecules that target key pathways involved in Asn metabolism, providing significant theoretical insights and potential drug candidates for the development of novel AD therapeutic strategies based on Asn metabolism modulation.

## 2. Materials and methods

### 2.1 Datasets and preprocessing

Microarray datasets associated with AD, including GSE5281, GSE29378, GSE36980, and GSE138260, ere accessed from the Gene Expression Omnibus (GEO) repository (https://www.ncbi.nlm.nih.gov/geo/). Corresponding platform annotation files were obtained to facilitate probe-to-gene symbol mapping. Clinical metadata including age, sex, and group assignment were also extracted. A list of AMGs was retrieved from the GeneCards human gene database (https://www.genecards.org/) by searching for the keyword ‘asparagine metabolism’. Genes with a relevance score > 6 and that were protein-coding were selected for analysis. A total of 1,294 AMGs were compiled for subsequent analyses. Data preprocessing was performed on each dataset, including background correction, log2 transformation, and quantile normalization of the raw expression values. Gene-level analysis was performed by mapping probe IDs to gene symbols, with the average expression value applied when multiple probes corresponded to a single gene. The data utilized in this study were publicly accessible, and no experimental work was carried out by the authors.

### 2.2 Identification of DEGs

Expression data were processed in R version 4.4.1 using the limma and sva packages. Batch effects were first assessed via principal component analysis (PCA), which allowed for the identification of significant inter-dataset variability. Genes with an adjusted *p*-value < 0.05 and absolute log2 fold change (|log2FC|) > 0.585 were considered significant. The DEGs from all datasets were intersected to identify common AD-related genes. DEG visualization was performed using the pheatmap package for heatmap generation and ggplot2 for volcano plot visualization. Venn diagram analyses were conducted using the VennDiagram package to intersect DEGs with the list of AMGs, ultimately identifying 39 candidate AMG-DEGs.

### 2.3 Functional enrichment analyses: GSEA, GO and KEGG

GSEA was performed using the clusterProfiler package in R, based on the reference gene set collections c5.go.Hs.symbols.gmt. The DEGs were ranked by their log2FC, and enrichment scores were computed for each gene set using the GSEA function. A threshold of an adjusted p-value < 0.05 was considered statistically significant.

GO and KEGG pathway enrichment analyses were performed using the “clusterProfiler” R package to explore cellular components (CC), molecular functions (MF), biological processes (BP), and signaling pathways associated with the intersecting AMG-DEGs. GO and KEGG enrichment analyses were conducted with the enrichGO and enrichKEGG functions, respectively, using the parameters: *p*-value < 0.05 and adjusted *p*-value < 0.05 as the significance thresholds.

### 2.4 Construction of the PPI network

To explore the interactions among the 39 AMG-DEGs, a PPI network was generated using the Retrieval of Interacting Genes (STRING) database (https://string-db.org/) for gene interaction retrieval, with a confidence score cutoff > 0.4. Unrelated nodes were removed to improve network clarity. The network was visualized using Cytoscape software (version 3.10.0), and hub modules were identified using the MCODE plugin with the following parameters: K-core = 2, node score cutoff = 0.2, degree cutoff = 2 and maximum depth = 100.

### 2.5 Hub gene identification and functional interaction analysis

The cytoHubba plugin in Cytoscape was employed to identify core AMG-DEGs using the Maximal Clique Centrality algorithm. Additionally, Gene Multiple Association Network Integration Algorithm (GeneMANIA) (http://www.genemania.org/) was used to construct gene co-expression networks, providing insight into potential functional relationships between hub genes.

### 2.6 Construction of the competing endogenous RNA (ceRNA) regulatory network

MiRNA-mRNA interactions targeting hub genes were predicted using multiple databases, including miRanda (http://www.microrna.org/), miRTarBase (http://mirtarbase.cuhk.edu.cn/), miRDB (http://mirdb.org/), and TargetScan (http://www.targetscan.org/). Long non-coding RNAs (lncRNAs)– miRNA interactions were obtained via SpongeScan (http://spongescan.rc.ufl.edu/). All predicted interactions were visualized in Cytoscape to generate the final ceRNA network, integrating lncRNA– miRNA and miRNA-mRNA regulatory axes.

### 2.7 Transcription factor regulatory network construction

Transcription factor (TF) regulating the identified hub genes were retrieved from the TRRUST database (https://www.grnpedia.org/trrust/). The TF-mRNA regulatory network was then visualized using Cytoscape to elucidate potential upstream regulatory mechanisms.

### 2.8 Drug enrichment analysis

Candidate drugs targeting the hub genes were identified from the Drug-Gene Interaction Database (DGIdb, https://www.dgidb.org/). Gene–compound association enrichment was performed using Drug Signatures Database (DSigDB) (https://maayanlab.cloud/DSigDB/), and results were visualized using the enrichplot package in R.

### 2.9 Molecular docking

Molecular docking was employed to assess the binding affinity between predicted small-molecule compounds and their target proteins. The top five candidate drugs—ranked by adjusted *p*-values from drug enrichment analysis—included DL-Mevalonic acid (MVA), Bathocuproine disulfonate (BCS), Phenethyl isothiocyanate (PEITC), MELAMINE, and CHLOROBENZENE. Corresponding protein structures were downloaded from the RCSB Protein Data Bank (PDB, https://www.rcsb.org/) for *CD44* (PDB ID: 1UUH), *CCL2* (PDB ID: 4USP), and *TUBB3* (PDB ID: 5IJ0). The protein structure of *NFKBIA* was predicted using the AlphaFold Protein Structure Database (https://www.alphafold.ebi.ac.uk/entry/P25963). 3D molecular docking simulations were performed with CB-Dock2 (https://cadd.labshare.cn/cb-dock2/php/index.php), and Vina score (binding energy ≤ −5.0 kcal/mol) were used to prioritize ligand–receptor interactions.

## 3. Results

### 3.1 Identification of AMG-DEGs in AD

To identify candidate genes associated with asparagine metabolism in AD, we analyzed four publicly available microarray datasets (GSE5281, GSE29378, GSE36980, and GSE138260). Clinical information including age, sex, and sample group distributions for each dataset is summarized in Table 1. PCA was performed to evaluate batch effects. Before correction, samples clustered separately by dataset, indicating strong batch effects (Fig 2A). After batch effect adjustment, the samples showed improved clustering consistency (Fig 2B), confirming effective normalization. Differential expression analysis revealed a total of 363 DEGs between AD and control samples, with 156 genes significantly upregulated and 207 downregulated (Fig 2C, S1 Fig). A Venn diagram analysis identified 39 genes overlapping between DEGs and AMGs (Fig 2D). The expression patterns of these 39 overlapping genes were visualized using a heatmap, which demonstrated distinct clustering between control and AD groups (Fig 2E). The complete list of these AMG-DEGs is provided in S1 Table.

**Fig 1.** Research design flow chart.

**Fig 2.** Expression profile analysis of overlapping genes between DEGs and AMGs. (A–B) PCA for batch effect assessment: (A) Before batch correction; (B) After batch correction. (C) Volcano plot displaying the results of DEGs analysis in AD, with red and blue dots representing upregulated and downregulated genes, respectively. (D) Venn diagram illustrating the intersection between DEGs and AMGs. (E) Heatmap of AMG-DEGs, with red and blue representing high and low expression levels, respectively.

**Table 1.** Microarrays datasets clinical characteristics.

| Dataset | GSE5281 |  | GSE29378 |  | GSE36980 |  | GSE138260 |  |
| --- | --- | --- | --- | --- | --- | --- | --- | --- |
| Groups | Control | AD | Control | AD | Control | AD | Control | AD |
| Number | 74 | 87 | 32 | 31 | 47 | 33 | 19 | 17 |
| Age | 73.30±18.3<br>5 | 79.53±6.8<br>4 | 81.65±6.7<br>7 | 76.64±8.9<br>6 | 78.09±9.3<br>5 | 91.70±5.8<br>3 | 64.31±17.1<br>5 | 79.82±9.3<br>4 |
| Male | 53 | 50 | 22 | 16 | 22 | 15 | 9 | 7 |
| **Female** | 21 | 37 | 10 | 15 | 25 | 18 | 9 | 10 |

### 3.2 Functional enrichment analyses

GSEA revealed that in control samples, energy production and neurotransmission-related pathways were enriched, including ATP synthesis coupled electron transport, GABA ergic synapse, inner mitochondrial membrane protein complex, mitochondrial protein containing complex and synaptic vesicle membrane (Fig 3A, S2 Table). In contrast, AD samples exhibited enrichment in pathways associated with positive regulation of vasculature development, regulation of epithelial cell differentiation, regulation of vasculature development, collagen containing extracellular matrix and growth factor binding (Fig 3B, S2 Table).

**Fig 3.** Functional enrichment analyses. (A-B) GSEA: (A) GO terms enriched in the control group; (B) GO terms enriched in the AD group; (C) GO enrichment bar plot showing the top 10 significantly enriched MF, CC, and BP. (D) GO enrichment bubble plot. (E) Circular plot of KEGG pathway enrichment analysis. (F) KEGG pathway enrichment bubble plot.

To further understand the biological roles of the 39 AMG-DEGs, we conducted GO and KEGG pathway enrichment analyses. The GO analysis revealed that these genes were significantly enriched in BP such as response to steroid hormones, regulation of neurogenesis, and nervous system development. For CC, the AMG-DEGs were found to be associated with the neuronal cell body, GABAergic synapses, and clathrin-coated vesicle membranes (Fig 3C-D, Table 2), emphasizing their involvement in maintaining neuronal function and synaptic communication. In terms of MF, the enriched terms included carbon-carbon lyase activity, carboxy-lyase activity, and pyridoxal phosphate (PLP) binding, all of which are essential for energy and neurotransmitter regulation (Fig 3C-D). KEGG pathway enrichment revealed that AMG-DEGs were significantly involved in pathways like GABAergic synapses (hsa04727), alanine, aspartate, and glutamate metabolism (hsa00250), and IL-17 signaling (hsa04657). Interestingly, pathways related to butanoate metabolism (hsa00650) and β-alanine metabolism (hsa00410) were found to be downregulated, while inflammation-related pathways, including IL-17 signaling and rheumatoid arthritis (hsa05323), were upregulated (Fig 3E-F).

**Table 2.** GO and KEGG enrichment analysis of AMG-DEGs.

| Term | ID | Description | GeneRatio | *p*.Value |
| --- | --- | --- | --- | --- |
| BP | GO:0048545 | response to steroid hormone | 8/39 | 3.67E-07 |
|  | GO:0043649 | dicarboxylic acid catabolic process | 3/39 | 5.42E-06 |
|  | GO:0050767 | regulation of neurogenesis | 7/39 | 1.39E-05 |
|  | GO:0009065 | glutamine family amino acid catabolic process | 3/39 | 2.05E-05 |
|  | GO:0016188 | synaptic vesicle maturation | 3/39 | 2.57E-05 |
| **CC** | GO:0098982 | GABA-ergic synapse | 5/39 | 4.50E-07 |
|  | GO:0030665 | clathrin-coated vesicle membrane | 5/39 | 7.11E-06 |
|  | GO:0043025 | neuronal cell body | 7/39 | 3.99E-05 |
|  | GO:0030662 | coated vesicle membrane | 5/39 | 4.79E-05 |
|  | GO:0030136 | clathrin-coated vesicle | 5/39 | 6.41E-05 |
| **MF** | GO:0016831 | carboxy-lyase activity | 3/39 | 4.94E-05 |
|  | GO:0016830 | carbon-carbon lyase activity | 3/39 | 1.68E-04 |
|  | GO:0030170 | pyridoxal phosphate binding | 3/39 | 2.21E-04 |
|  | GO:0070279 | vitamin B6 binding | 3/39 | 2.33E-04 |
|  | GO:0008503 | benzodiazepine receptor activity | 2/39 | 2.35E-04 |
| **KEGG** | hsa04727 | GABAergic synapse | 4/34 | 4.02E-04 |
|  | hsa00250 | Alanine, aspartate and glutamate metabolism | 3/34 | 4.08E-04 |
|  | hsa00430 | Taurine and hypotaurine metabolism | 2/34 | 2.01E-03 |
|  | hsa05120 | Epithelial cell signaling in *Helicobacter pylori* infection | 3/34 | 2.74E-03 |
|  | hsa00650 | Butanoate metabolism | 2/34 | 5.07E-03 |

### 3.3 Construction of PPI network and module analysis

A PPI network was constructed for the 39 AMG-DEGs using the STRING database with a confidence score threshold of >0.4. The network was visualized in Cytoscape, which revealed a network consisting of 34 nodes and 78 edges, with 15 upregulated genes highlighted in pink and 19 downregulated genes in green (Fig 4A). The MCODE plugin was used to identify key network modules, which revealed two significant clusters. Module 1 contained 7 hub genes and 21 interactions, which were primarily enriched in genes related to GABAergic neurotransmission, such as *SST*, *GAD1*, and *GAD2*. This suggests a central role for these genes in neurotransmitter regulation (Fig 4B). Module 2 was composed of immune-related genes, including *CCL2*, *CXCR4*, and *CD44*, which are implicated in immune response and inflammatory signaling (Fig 4C). These results highlight the involvement of AMG-DEGs in both neurotransmission and immune regulation, suggesting that disrupted pathways in AD may affect both neural and inflammatory processes.

**Fig 4.** PPI network of AMG-DEGs and functional subclusters. (A) PPI network constructed from AMG-DEGs. (B-C) Subclusters extracted from the PPI network. Red nodes indicate upregulated genes, green nodes indicate downregulated genes.

### 3.4 Identification and characterization of core AMG-DEGs

Using the UpSetR package and 10 centrality metrics in the cytoHubba plugin, we identified the top 20 genes per metric and obtained the intersection, resulting in 7 consensus hub genes: *GFAP*, *CCL2*, *NFKBIA*, *TUBB3*, *GAD2*, *CD44*, and *HPRT1* (Fig 5A, S3 Table). Table 3 summarizes their full names and functional annotations. GeneMANIA analysis of these hub genes revealed a complex co-expression network dominated by co-expression interactions (80.85%), with co-localization relationships accounting for the remaining 19.15% (Fig 5B). Key pathways identified as significantly enriched included responses to bacterial molecules and lipopolysaccharides, regulation of leukocyte cell-cell adhesion, cellular reactions to biotic stimuli, as well as negative regulation of apoptotic signaling, among others.

**Fig 5.** Identification of key hub genes among AMG - DEGs. (A) UpSet plot showing the intersection of seven hub genes identified by 10 different computational algorithms. (B) GeneMANIA analysis of core AMG-DEGs and their co-expressed genes.

**Table 3.**
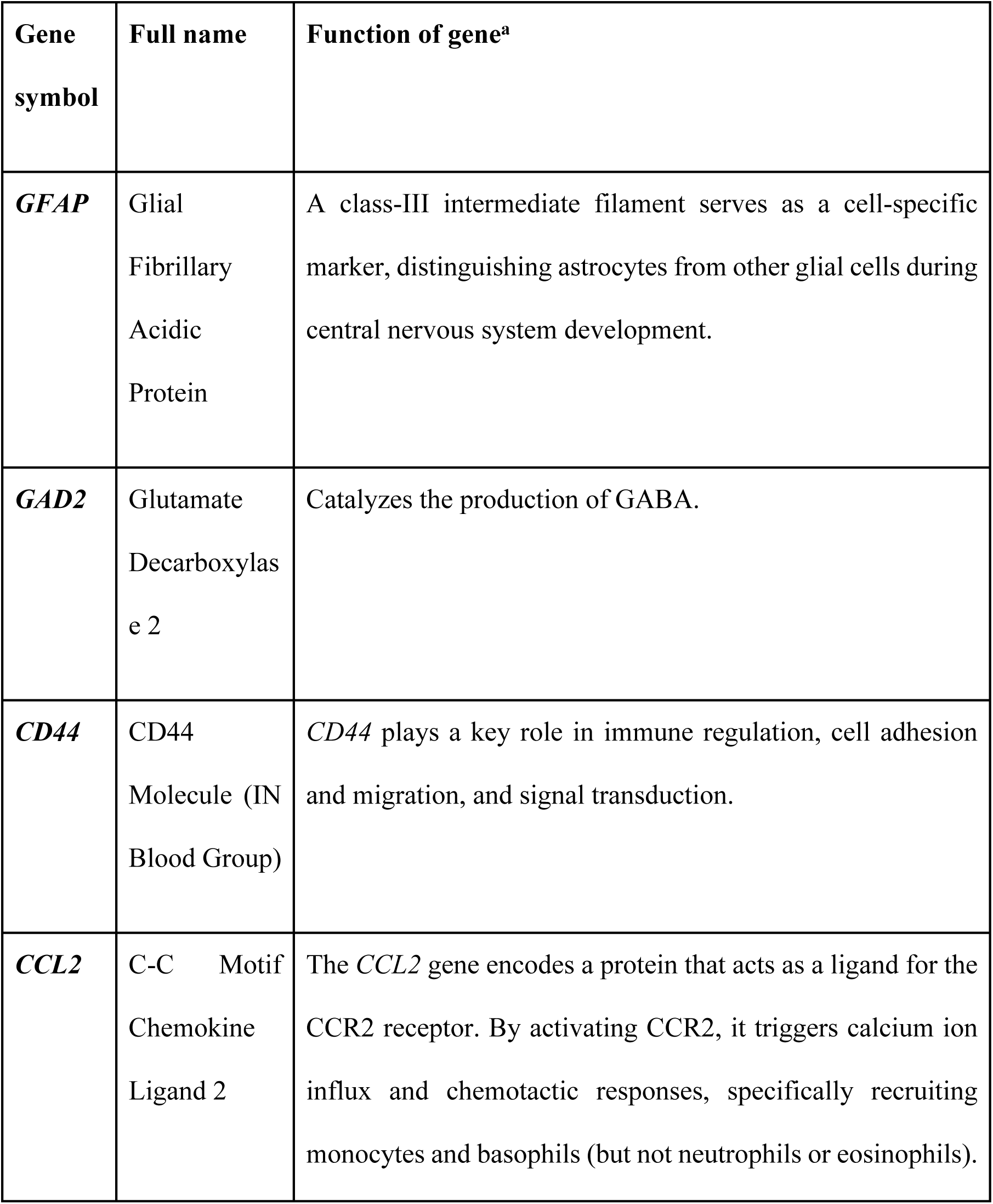
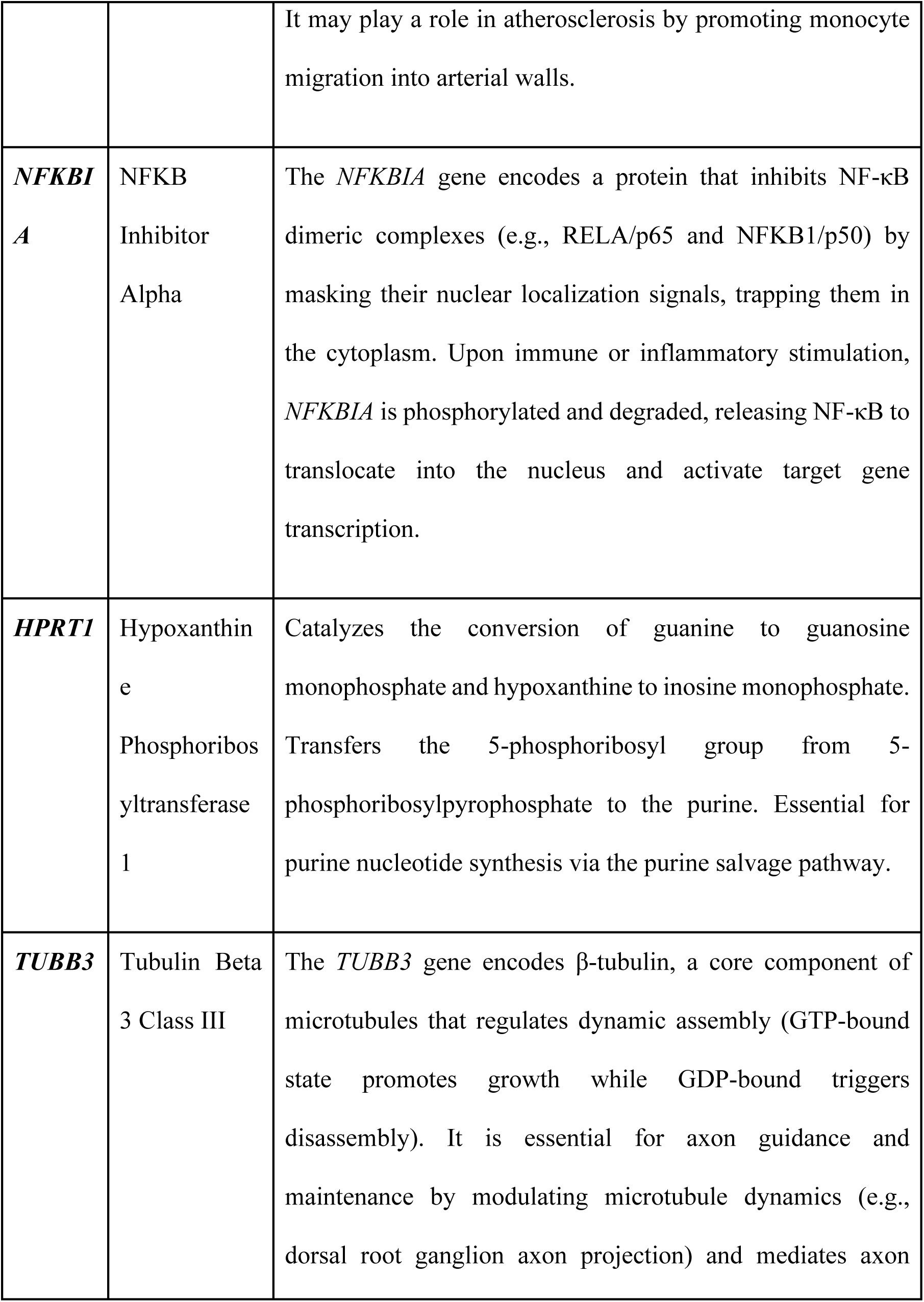
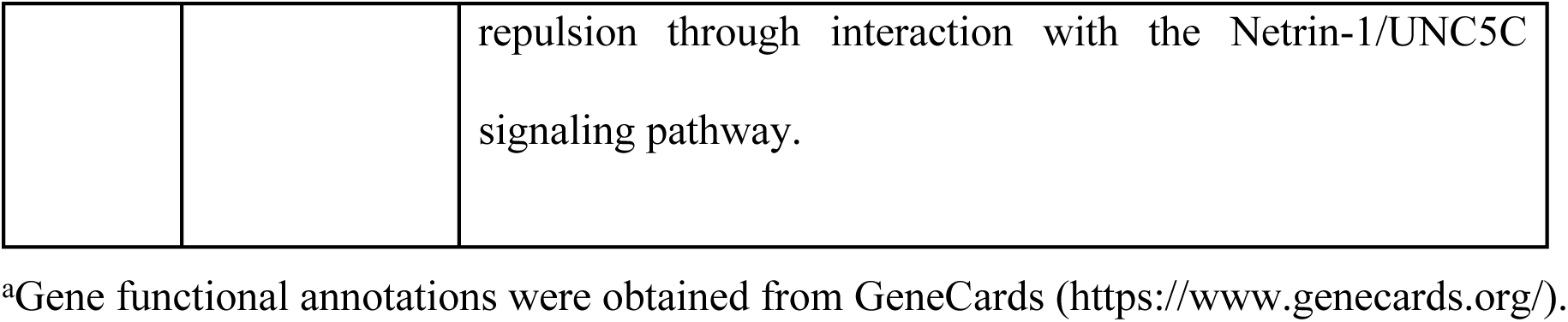
The details of the core AMG-DEGs.

### 3.5 Construction of a ceRNA regulatory network

To further elucidate the post-transcriptional regulatory mechanisms associated with AD, we constructed a ceRNA regulatory network (Fig 6A). In this network, *HPRT1*, *CD44*, and *CCL2* emerged as central nodes, suggesting their potential roles in asparagine metabolism-related ceRNA regulation. lncRNAs including RP4-539M6.22, CITF22-1A6.3, LA16c-306A4.2, and SNHG14 were predicted to regulate the expression of *HPRT1* mRNA by competitively binding to hsa-miR-130a-3p. Similarly, CTC-459F4.1 was found to potentially modulate *HPRT1* through competition for hsa-miR-576-5p. In the regulation of *CCL2*, lncRNAs LINC01043, GNG12-AS1, and RP3-470B24.5 were identified as ceRNAs via their interaction with hsa-miR-1-3p. Moreover, lncRNAs GS1-251I9.3, CTC-457E21.1, and RP11-486P11.1 were predicted to regulate *CD44* expression through competition for hsa-miR-130b-5p. These lncRNA-miRNA-mRNA regulatory axes may play critical roles in the pathogenesis of AD.

**Fig 6.** Regulatory networks of ceRNAs and transcription factors. (A) ceRNA regulatory network: green hexagons represent miRNAs, blue rhomboids represent lncRNAs, and red circles represent mRNAs. (B) TF regulatory network: green rectangles represent TF, red ellipses represent target genes.

### 3.6 Construction of a transcription factor regulatory network

A transcriptional regulatory network consisting of 9 nodes and 12 interactions was established based on known transcription factor (TF)-target relationships. Our analysis revealed that *CCL2*, a key pro-inflammatory cytokine, is regulated by multiple TFs, including *NFKB1*, *STAT3*, *SP1*, *NFIC*, and *RELA*. Likewise, the expression of *GFAP* is modulated by *NFKB1*, *STAT3*, *NFIC*, and *RELA*. *NFKBIA* was found to be jointly regulated by *NFKB1* and *RELA*, while *CD44* is transcriptionally controlled by *SP1*. These results highlight *NFKB1*, *STAT3*, *SP1*, *NFIC*, and *RELA* as core transcriptional regulators of AMG-DEGs in AD (Fig 6B).

### 3.7 Identification of potential therapeutic compounds

To explore therapeutic strategies targeting key AMG-DEGs, we conducted drug enrichment analysis. The top candidate compounds with the highest statistical significance included MVA, BCS, PEITC, MELAMINE, and CHLOROBENZENE (Fig 7A). A drug-gene interaction network was constructed to further elucidate the molecular mechanisms underlying these compounds (Fig 7B). The network revealed potential interactions between the identified compounds and the hub genes, suggesting that these molecules may exert therapeutic effects by modulating inflammation-or neurofunction-related pathways involved in AD pathogenesis.

**Fig 7.** Drug enrichment analysis and gene-drug interaction network. (A) Bar plot of drug enrichment results. (B) Drug-gene interaction network illustrating the potential associations between identified drugs and target genes.

### 3.8 Molecular docking analysis

To validate the therapeutic potential of the candidate compounds, molecular docking analysis was performed using CB-Dock, and the binding affinities were evaluated using Vina scores (Table 4, Fig 8). Among the compounds, BCS exhibited the strongest binding affinity, particularly with CD44, showing a Vina score of −9.3 kcal/mol (Fig 8C), outperforming MVA (−5.7 kcal/mol, Fig 8A). Additionally, BCS demonstrated high binding affinities with CCL2 and NFKBIA (Vina scores: −7.6 and −7.1 kcal/mol, respectively; Fig 8B, D), supporting its potential as a multi-target modulator. PEITC showed a favorable binding affinity to TUBB3 (−6.2 kcal/mol) with a relatively large cavity volume (1608 Å³, Fig 8E), although its interaction with CCL2 and NFKBIA was relatively weak (Vina scores: −4.2 and −4.4 kcal/mol, respectively; S2 Fig). In contrast, MELAMINE and CHLOROBENZENE exhibited poor binding capacities across all docking analyses, with the highest Vina scores of −4.2 and −3.8 kcal/mol, respectively, suggesting limited therapeutic potential (S2 Fig).

**Fig 8.** Molecular docking results of small-molecule compounds with target proteins. (A) Molecular docking of DL-Mevalonic acid with CD44 protein. (B) Docking result of Bathocuproine disulfonate with NFKBIA protein. (C) Docking result of Bathocuproine disulfonate with CD44 protein. (D) Docking result of Bathocuproine disulfonate with CCL2 protein. (E) Docking result of Phenethyl isothiocyanate with TUBB3 protein.

**Table 4.** Binding affinity and pocket volume of drug-protein complexes identified by molecular docking.

| Drug | Protein | Vina score<br>(kcal/mol) | Cavity volume<br>(Å<sup>3</sup>) |
| --- | --- | --- | --- |
| DL-Mevalonic acid | CD44 | -5.7 | 121 |
| DL-Mevalonic acid | CCL2 | -4.1 | 82 |
| **DL-Mevalonic acid** | NFKBIA | -3.7 | 285 |
| **Bathocuproine disulfonate** | CD44 | -9.3 | 308 |
| **Bathocuproine disulfonate** | CCL2 | -7.6 | 102 |
| **Bathocuproine disulfonate** | NFKBIA | -7.1 | 72 |
| **Phenethyl isothiocyanate** | CCL2 | -4.2 | 107 |
| **Phenethyl isothiocyanate** | NFKBIA | -4.4 | 72 |
| **Phenethyl isothiocyanate** | TUBB3 | -6.2 | 1608 |
| **MELAMINE** | CCL2 | -4.2 | 107 |
| **MELAMINE** | NFKBIA | -4.2 | 285 |
| **CHLOROBENZENE** | CCL2 | -3.5 | 102 |
| **CHLOROBENZENE** | NFKBIA | -3.8 | 72 |

## Discussion

AD remains one of the most widespread neurodegenerative conditions globally, particularly affecting the aging population. Despite recent advances in AD diagnosis and treatment, the disease remains incurable, with current therapeutic strategies primarily aimed at alleviating symptoms rather than halting disease progression(23). Identifying new molecular targets involved in the core processes of AD is increasingly important. This study offers fresh insights into how Asn metabolism may contribute to AD progression. Our results emphasize the role of key genes and regulatory mechanisms that impact neuroinflammation, synaptic function, and energy balance in AD.

GSEA in this study revealed impairments in energy metabolism, synaptic dysfunction, and vascular dysfunction of AD, which aligns with previous reports(24, 25). GO analysis further supported this, indicating that genes involved in Asn metabolism are primarily engaged in neurogenesis, nervous system development, and synaptic plasticity—processes tightly linked to cognitive decline in AD(26, 27). In particular, the CC enrichment results showed significant AMG-DEGs localization in neuronal cell bodies, GABAergic synapses, and clathrin-coated vesicle membranes, indicating that asparagine metabolism may impact synaptic signaling and vesicle transport. Previous studies have demonstrated that GABAergic synapses are essential for inhibitory neural regulation(28), and disruptions in this system have been associated with synaptic abnormalities in AD and other neurodegenerative diseases(29, 30). Interestingly, MF enrichment analysis revealed a significant association with carbon-carbon lyase activity and pyridoxal phosphate binding. Pyridoxal phosphate is a crucial cofactor for neurotransmitter biosynthesis(31), indicating that alterations in Asn metabolism may disrupt neurotransmitter balance in AD. KEGG pathway analysis revealed a downregulation in butyrate and β-alanine metabolism pathways, while inflammation-related pathways, such as the IL-17 signaling pathway, were upregulated. This inverse regulation between metabolism and inflammation suggests an imbalance in the immune-metabolic axis in AD. Although previous studies demonstrated the dynamic interplay between neuroinflammation and disturbances in lipid, glucose, and amino acid metabolism in AD(32, 33), our study identified asparagine metabolism as a previously unrecognized key node within the immune-metabolic axis. This metabolite-specific mechanism not only supports the concept of the immune-metabolic axis but also provides further evidence for how discrete metabolic dysregulation can modulate neuroinflammation, thus refining the theoretical framework of immune-metabolism in AD.

Among the seven hub genes identified in our study, *GFAP*, *CD44*, *CCL2*, and *NFKBIA* were upregulated, while *GAD2*, *TUBB3*, and *HPRT1* were downregulated. These genes play pivotal roles in inflammation, synaptic stability, and neurotransmitter regulation, which are critical to AD pathophysiology. The upregulation of neuroinflammatory markers like *GFAP*(34), *CD44*(35), and *CCL2*(36) in AD corroborates findings from prior studies linking chronic inflammation to disease progression(35, 37–39). Upregulation of *CCL2*(40) and *NFKBIA*(41) is associated with the IL-17 signaling pathway, further validating the results from the KEGG pathway analysis. Furthermore, the downregulation of *GAD2*, which encodes glutamate decarboxylase responsible for GABA synthesis, is consistent with the known imbalance in excitatory-inhibitory neurotransmission in AD(42, 43). Our findings show that both *TUBB3* and *HPRT1* are downregulated in AD. Research has shown that downregulation of *TUBB3* may destabilize microtubules, leading to Tau hyperphosphorylation and neuronal structural damage in AD(44). Additionally, consistent with extensive literature on energy metabolism imbalances in AD(45, 46), our results suggest that the downregulation of *HPRT1* could impair purine metabolism, leading to insufficient energy supply and mitochondrial dysfunction in neurons.

The regulatory networks constructed in this study also highlighted the complexity of post-transcriptional regulation in AD. The ceRNA network analysis suggested that multiple lncRNAs and miRNAs regulate hub genes such as *HPRT1*, *CD44*, and *CCL2*. These interactions may modulate the inflammatory and energy metabolism in AD. Although previous studies have shown that under ischemic conditions, SNHG14 exacerbates neuronal damage through the miR-182-5p/*BINP3* signaling axis(47), our study identifies a novel mechanism in AD, wherein SNHG14 modulates the miR-130a-3p/*HPRT1* pathway to influence purine metabolism. This differential regulatory pattern suggests that SNHG14 may act as a ‘metabolic switch’, maintaining energy homeostasis by regulating mitochondrial autophagy during acute injury, while modulating energy supply through the purine metabolism pathway in chronic neurodegenerative conditions. The TF network revealed the involvement of *NFKB1*, *STAT3*, and *RELA* in regulating these pathways, emphasizing their potential as therapeutic targets in modulating the immune response and synaptic plasticity. Our analysis revealed that *CD44* is uniquely regulated by the transcription factor *SP1*, suggesting that *CD44* may influence inflammatory responses through an independent pathway(48). Notably, *NFKBIA*, a negative regulator of NF-κB signaling, appears to act as a compensatory mechanism to suppress excessive inflammation. However, this inhibition may be insufficient to counterbalance the chronic neuroinflammation observed in AD progression(49).

The potential therapeutic compounds identified through drug enrichment and molecular docking analysis, including BCS, MVA, and PEITC, show promising results in targeting key genes and pathways involved in AD. Molecular docking results revealed that Bisulfate-derived compound BCS exhibits a strong binding affinity to CD44, CCL2, and NFKBIA proteins, suggesting that BCS may alleviate the progression of AD through the inhibition of inflammatory pathways. Previous studies have indicated that excessive copper in Alzheimer’s disease may exacerbate Aβ aggregation and oxidative stress(50). BCS, a copper chelator, has shown potential therapeutic value by restoring metal ion balance and may therefore provide beneficial effects in the context of AD(51, 52). MVA, a key intermediate in cholesterol biosynthesis, was significantly enriched in our analysis. Cholesterol metabolism dysregulation has been implicated in AD pathogenesis(53, 54). Statins, which target this pathway, have shown promise in AD prevention and treatment by reducing Aβ accumulation, suppressing inflammation, enhancing vascular function, and modulating Tau phosphorylation(55). Nonetheless, the optimal therapeutic window and specific statin formulations for AD remain under debate. Intriguingly, we found that MVA exhibits strong binding affinity to CD44 proteins, a finding not widely reported in previous studies. This suggests a potential link between MVA and CD44 signaling in AD, warranting further investigation. PEITC is a naturally occurring compound known for its antioxidant and anti-inflammatory properties(56). Our docking analysis indicated significant binding affinity between PEITC and TUBB3 proteins, suggesting that PEITC may help stabilize microtubule structures and mitigate neuronal damage associated with AD. Although the binding affinity of PEITC to CCL2 and NFKBIA proteins was relatively weaker, a considerable body of evidence suggests that PEITC not only inhibits Aβ aggregation in AD(57), but also exerts its therapeutic effects through antioxidant and anti-inflammatory mechanisms, thereby slowing the disease’s progression(58, 59). However, despite promising preclinical findings, clinical studies investigating the efficacy of BCS, MVA, and PEITC in AD treatment are currently lacking, and further validation of their practical effects is required.

This study comprehensively elucidated the potential involvement of asparagine metabolism in AD pathogenesis and identified several key molecular targets. Nevertheless, some limitations should be noted. First, our findings are primarily based on public datasets and bioinformatics predictions, requiring further validation through experiments. Second, the clinical translational potential of our molecular docking results needs to be further assessed. Future studies integrating single-cell sequencing, metabolomics, and experimental validation are warranted to clarify the role of asparagine metabolism in AD and evaluate its feasibility as a therapeutic target.

### Conclusion

In conclusion, this study sheds light on the role of asparagine metabolism in AD. We identified key genes associated with neuroinflammation, synaptic function, and energy metabolism, suggesting they could be potential targets for therapy. Our analysis of regulatory networks also uncovered intricate interactions between miRNAs, lncRNAs, and transcription factors that influence these genes. Moreover, drug screening and molecular docking revealed several promising compounds that may offer therapeutic benefits for AD. These findings indicate that modulating asparagine metabolism could be a new strategy for AD treatment. However, further research is needed to validate these results experimentally.

## Acknowledgments

We gratefully acknowledge all contributors for their valuable participation in this study.

## Funding Statement

This study was supported by the Yunnan Science and Technology Program (Grant/Award Number: 202401AT070176).

## Abbreviations

Aβ, β-amyloid; AD, Alzheimer’s disease; AEP, asparagine endopeptidase; AMGs, asparagine metabolism-related genes; AMG - DEGs, asparagine metabolism- differentially expressed genes; Asn, asparagine; BCS, Bathocuproine disulfonate; Betweenness, Betweenness Centrality; BottleNeck, BottleNeck Algorithm; BP, biological process; CC, cellular component; ceRNA, competing endogenous RNA; Closeness, Closeness Centrality; DEG, differentially expressed gene; Degree, Degree Centrality; DGIdb, Drug-Gene Interaction Database; DMNC, Density of Maximum Neighborhood Component; DSigDB, Drug Signatures Database; EcCentricity, Eccentricity Centrality; GeneMANIA, Gene Multiple Association Network Integration Algorithm; GO, Gene Ontology; KEGG, Kyoto Encyclopedia of Genes and Genomes; lncRNAs, Long non-coding RNAs; MCC, Maximal Clique Centrality; MF, molecular function; MNC, Maximum Neighborhood Component; MVA, DL-Mevalonic acid; PCA, principal component analysis; PEITC, Phenethyl isothiocyanate; PPI, protein-protein interaction; Radiality, Radiality Centrality; Stress, Stress Centrality; STRING, Search Tool for the Retrieval of Interacting Genes; TCA, tricarboxylic acid; TF, Transcription factor.

## Notes

### Competing Interest Statement

The authors have declared no competing interest.

## References

1. Niu Q, Li D, Zhang J, Piao Z, Xu B, Xi Y, et al. The new perspective of Alzheimer’s Disease Research: Mechanism and therapeutic strategy of neuronal senescence. Ageing Res Rev. 2024;102:102593. Epub 2024/11/21. doi: 10.1016/j.arr.2024.102593. PubMed PMID: 39566741.

2. Tenchov R, Sasso JM, Zhou QA. Alzheimer’s Disease: Exploring the Landscape of Cognitive Decline. ACS Chem Neurosci. 2024;15(21):3800–27. Epub 2024/10/11. doi: 10.1021/acschemneuro.4c00339. PubMed PMID: 39392435; PubMed Central PMCID: PMC11587518.

3. Azargoonjahromi A. The duality of amyloid-β: its role in normal and Alzheimer’s disease states. Mol Brain. 2024;17(1):44. Epub 2024/07/18. doi: 10.1186/s13041-024-01118-1. PubMed PMID: 39020435; PubMed Central PMCID: PMC11256416.

4. Wang S, Jiang Y, Yang A, Meng F, Zhang J. The Expanding Burden of Neurodegenerative Diseases: An Unmet Medical and Social Need. Aging Dis. 2024. Epub 2024/11/21. doi: 10.14336/ad.2024.1071. PubMed PMID: 39571158.

5. Zhang Y, Li Y, Ma L. Recent advances in research on Alzheimer’s disease in China. J Clin Neurosci. 2020;81:43–6. Epub 2020/11/24. doi: 10.1016/j.jocn.2020.09.018. PubMed PMID: 33222956.

6. Peng Y, Gao P, Shi L, Chen L, Liu J, Long J. Central and Peripheral Metabolic Defects Contribute to the Pathogenesis of Alzheimer’s Disease: Targeting Mitochondria for Diagnosis and Prevention. Antioxid Redox Signal. 2020;32(16):1188–236. Epub 2020/02/14. doi: 10.1089/ars.2019.7763. PubMed PMID: 32050773; PubMed Central PMCID: PMC7196371.

7. Xu L, Liu R, Qin Y, Wang T. Brain metabolism in Alzheimer’s disease: biological mechanisms of exercise. Transl Neurodegener. 2023;12(1):33. Epub 2023/06/27. doi: 10.1186/s40035-023-00364-y. PubMed PMID: 37365651; PubMed Central PMCID: PMC10294518.

8. Ezkurdia A, Ramírez MJ, Solas M. Metabolic Syndrome as a Risk Factor for Alzheimer’s Disease: A Focus on Insulin Resistance. Int J Mol Sci. 2023;24(5). Epub 2023/03/12. doi: 10.3390/ijms24054354. PubMed PMID: 36901787; PubMed Central PMCID: PMC10001958.

9. Yue C, Fu Y, Zhao Y, Ou Y, Sun Y, Tan L. Association between Alzheimer’s disease and metabolic syndrome: Unveiling the role of dyslipidemia mechanisms. Brain Network Disorders. 2025;1(1):21–7. doi: 10.1016/j.bnd.2024.10.006.

10. Lauretti E, Dabrowski K, Praticò D. The neurobiology of non-coding RNAs and Alzheimer’s disease pathogenesis: Pathways, mechanisms and translational opportunities. Ageing Research Reviews. 2021;71:101425. doi: 10.1016/j.arr.2021.101425.

11. Chang H-C, Tsai C-Y, Hsu C-L, Tai T-S, Cheng M-L, Chuang Y-M, et al. Asparagine deprivation enhances T cell antitumour response in patients via ROS-mediated metabolic and signal adaptations. Nature Metabolism. 2025. doi: 10.1038/s42255-025-01245-6.

12. Lomelino CL, Andring JT, McKenna R, Kilberg MS. Asparagine synthetase: Function, structure, and role in disease. J Biol Chem. 2017;292(49):19952–8. Epub 2017/11/01. doi: 10.1074/jbc.R117.819060. PubMed PMID: 29084849; PubMed Central PMCID: PMC5723983.

13. Novotny BC, Fernandez MV, Wang C, Budde JP, Bergmann K, Eteleeb AM, et al. Metabolomic and lipidomic signatures in autosomal dominant and late-onset Alzheimer’s disease brains. Alzheimers Dement. 2023;19(5):1785–99. Epub 2022/10/18. doi: 10.1002/alz.12800. PubMed PMID: 36251323; PubMed Central PMCID: PMC10106526.

14. Chatterjee P, Cheong YJ, Bhatnagar A, Goozee K, Wu Y, McKay M, et al. Plasma metabolites associated with biomarker evidence of neurodegeneration in cognitively normal older adults. J Neurochem. 2021;159(2):389–402. Epub 2020/07/18. doi: 10.1111/jnc.15128. PubMed PMID: 32679614.

15. Alkan HF, Bogner-Strauss JG. Maintaining cytosolic aspartate levels is a major function of the TCA cycle in proliferating cells. Mol Cell Oncol. 2019;6(5):e1536843. Epub 2019/09/19. doi: 10.1080/23723556.2018.1536843. PubMed PMID: 31528687; PubMed Central PMCID: PMC6736317.

16. González-Domínguez R, García-Barrera T, Gómez-Ariza JL. Metabolite profiling for the identification of altered metabolic pathways in Alzheimer’s disease. J Pharm Biomed Anal. 2015;107:75–81. Epub 2015/01/13. doi: 10.1016/j.jpba.2014.10.010. PubMed PMID: 25575172.

17. Bradberry MM, Peters-Clarke TM, Shishkova E, Chapman ER, Coon JJ. N-glycoproteomics of brain synapses and synaptic vesicles. Cell Rep. 2023;42(4):112368. Epub 2023/04/11. doi: 10.1016/j.celrep.2023.112368. PubMed PMID: 37036808; PubMed Central PMCID: PMC10560701.

18. Shimonaka S, Matsumoto SE, Elahi M, Ishiguro K, Hasegawa M, Hattori N, et al. Asparagine residue 368 is involved in Alzheimer’s disease tau strain-specific aggregation. J Biol Chem. 2020;295(41):13996–4014. Epub 2020/08/08. doi: 10.1074/jbc.RA120.013271. PubMed PMID: 32759167; PubMed Central PMCID: PMC7549045.

19. Tang TMS, Luk LYP. Asparaginyl endopeptidases: enzymology, applications and limitations. Org Biomol Chem. 2021;19(23):5048–62. Epub 2021/05/27. doi: 10.1039/d1ob00608h. PubMed PMID: 34037066; PubMed Central PMCID: PMC8209628.

20. Song M. The Asparaginyl Endopeptidase Legumain: An Emerging Therapeutic Target and Potential Biomarker for Alzheimer’s Disease. Int J Mol Sci. 2022;23(18). Epub 2022/09/24. doi: 10.3390/ijms231810223. PubMed PMID: 36142134; PubMed Central PMCID: PMC9499314.

21. Behrendt A, Bichmann M, Ercan-Herbst E, Haberkant P, Schöndorf DC, Wolf M, et al. Asparagine endopeptidase cleaves tau at N167 after uptake into microglia. Neurobiology of Disease. 2019;130:104518. doi: 10.1016/j.nbd.2019.104518.

22. Meng X, Li B, Wang M, Zheng W, Ye K. Development of asparagine endopeptidase inhibitors for treating neurodegenerative diseases. Trends in Molecular Medicine. 2025;31(4):359–72. doi: 10.1016/j.molmed.2025.01.009.

23. Ogos M, Stary D, Bajda M. Recent Advances in the Search for Effective Anti-Alzheimer’s Drugs. Int J Mol Sci. 2024;26(1). Epub 2025/01/11. doi: 10.3390/ijms26010157. PubMed PMID: 39796014; PubMed Central PMCID: PMC11720639.

24. Wilson DM, 3rd, Cookson MR, Van Den Bosch L, Zetterberg H, Holtzman DM, Dewachter I. Hallmarks of neurodegenerative diseases. Cell. 2023;186(4):693–714. Epub 2023/02/22. doi: 10.1016/j.cell.2022.12.032. PubMed PMID: 36803602.

25. Yuan Y, Zhao G, Zhao Y. Dysregulation of energy metabolism in Alzheimer’s disease. J Neurol. 2024;272(1):2. Epub 2024/12/02. doi: 10.1007/s00415-024-12800-8. PubMed PMID: 39621206; PubMed Central PMCID: PMC11611936.

26. Zhang X, Wei X, Mei Y, Wang D, Wang J, Zhang Y, et al. Modulating adult neurogenesis affects synaptic plasticity and cognitive functions in mouse models of Alzheimer’s disease. Stem Cell Reports. 2021;16(12):3005–19. Epub 2021/12/04. doi: 10.1016/j.stemcr.2021.11.003. PubMed PMID: 34861165; PubMed Central PMCID: PMC8693766.

27. Peng L, Bestard-Lorigados I, Song W. The synapse as a treatment avenue for Alzheimer’s Disease. Mol Psychiatry. 2022;27(7):2940–9. Epub 2022/04/22. doi: 10.1038/s41380-022-01565-z. PubMed PMID: 35444256.

28. Lepeta K, Lourenco MV, Schweitzer BC, Martino Adami PV, Banerjee P, Catuara-Solarz S, et al. Synaptopathies: synaptic dysfunction in neurological disorders - A review from students to students. J Neurochem. 2016;138(6):785–805. Epub 2016/06/23. doi: 10.1111/jnc.13713. PubMed PMID: 27333343; PubMed Central PMCID: PMC5095804.

29. Krueger-Burg D. Understanding GABAergic synapse diversity and its implications for GABAergic pharmacotherapy. Trends Neurosci. 2025;48(1):47–61. Epub 2025/01/09. doi: 10.1016/j.tins.2024.11.007. PubMed PMID: 39779392.

30. Tang X, Jaenisch R, Sur M. The role of GABAergic signalling in neurodevelopmental disorders. Nat Rev Neurosci. 2021;22(5):290–307. Epub 2021/03/28. doi: 10.1038/s41583-021-00443-x. PubMed PMID: 33772226; PubMed Central PMCID: PMC9001156.

31. Bisello G, Longo C, Rossignoli G, Phillips RS, Bertoldi M. Oxygen reactivity with pyridoxal 5’-phosphate enzymes: biochemical implications and functional relevance. Amino Acids. 2020;52(8):1089–105. Epub 2020/08/28. doi: 10.1007/s00726-020-02885-6. PubMed PMID: 32844248; PubMed Central PMCID: PMC7497351.

32. Jung ES, Choi H, Mook-Jung I. Decoding microglial immunometabolism: a new frontier in Alzheimer’s disease research. Mol Neurodegener. 2025;20(1):37. Epub 2025/03/28. doi: 10.1186/s13024-025-00825-0. PubMed PMID: 40149001; PubMed Central PMCID: PMC11948825.

33. Chen H, Guo Z, Sun Y, Dai X. The immunometabolic reprogramming of microglia in Alzheimer’s disease. Neurochemistry International. 2023;171:105614. doi: 10.1016/j.neuint.2023.105614.

34. Li D, Liu X, Liu T, Liu H, Tong L, Jia S, et al. Neurochemical regulation of the expression and function of glial fibrillary acidic protein in astrocytes. Glia. 2020;68(5):878–97. Epub 2019/10/19. doi: 10.1002/glia.23734. PubMed PMID: 31626364.

35. Al-Dalahmah O, Sosunov AA, Sun Y, Liu Y, Madden N, Connolly ES, et al. The Matrix Receptor CD44 Is Present in Astrocytes throughout the Human Central Nervous System and Accumulates in Hypoxia and Seizures. Cells. 2024;13(2). Epub 2024/01/22. doi: 10.3390/cells13020129. PubMed PMID: 38247821; PubMed Central PMCID: PMC10814649.

36. Guo S, Zhang Q, Guo Y, Yin X, Zhang P, Mao T, et al. The role and therapeutic targeting of the CCL2/CCR2 signaling axis in inflammatory and fibrotic diseases. Front Immunol. 2024;15:1497026. Epub 2025/01/24. doi: 10.3389/fimmu.2024.1497026. PubMed PMID: 39850880; PubMed Central PMCID: PMC11754255.

37. Roveta F, Bonino L, Piella EM, Rainero I, Rubino E. Neuroinflammatory Biomarkers in Alzheimer’s Disease: From Pathophysiology to Clinical Implications. Int J Mol Sci. 2024;25(22). Epub 2024/11/27. doi: 10.3390/ijms252211941. PubMed PMID: 39596011; PubMed Central PMCID: PMC11593837.

38. Kölliker-Frers R, Udovin L, Otero-Losada M, Kobiec T, Herrera MI, Palacios J, et al. Neuroinflammation: An Integrating Overview of Reactive-Neuroimmune Cell Interactions in Health and Disease. Mediators Inflamm. 2021;2021:9999146. Epub 2021/06/24. doi: 10.1155/2021/9999146. PubMed PMID: 34158806; PubMed Central PMCID: PMC8187052.

39. Hu D, Mo X, Jihang L, Huang C, Xie H, Jin L. Novel diagnostic biomarkers of oxidative stress, immunological characterization and experimental validation in Alzheimer’s disease. Aging (Albany NY). 2023;15(19):10389–406. Epub 2023/10/06. doi: 10.18632/aging.205084. PubMed PMID: 37801482; PubMed Central PMCID: PMC10599743.

40. Ruiz de Morales JMG, Puig L, Daudén E, Cañete JD, Pablos JL, Martín AO, et al. Critical role of interleukin (IL)-17 in inflammatory and immune disorders: An updated review of the evidence focusing in controversies. Autoimmunity Reviews. 2020;19(1):102429. doi: 10.1016/j.autrev.2019.102429.

41. Liu Y, Meng Y, Zhou C, Yan J, Guo C, Dong W. Activation of the IL-17/TRAF6/NF-κB pathway is implicated in Aβ-induced neurotoxicity. BMC Neurosci. 2023;24(1):14. Epub 2023/02/25. doi: 10.1186/s12868-023-00782-8. PubMed PMID: 36823558; PubMed Central PMCID: PMC9951515.

42. Chiu CQ, Barberis A, Higley MJ. Preserving the balance: diverse forms of long-term GABAergic synaptic plasticity. Nat Rev Neurosci. 2019;20(5):272–81. Epub 2019/03/07. doi: 10.1038/s41583-019-0141-5. PubMed PMID: 30837689.

43. Koh W, Kwak H, Cheong E, Lee CJ. GABA tone regulation and its cognitive functions in the brain. Nat Rev Neurosci. 2023;24(9):523–39. Epub 2023/07/27. doi: 10.1038/s41583-023-00724-7. PubMed PMID: 37495761.

44. Radwitz J, Hausrat TJ, Heisler FF, Janiesch PC, Pechmann Y, Rübhausen M, et al. Tubb3 expression levels are sensitive to neuronal activity changes and determine microtubule growth and kinesin-mediated transport. Cell Mol Life Sci. 2022;79(11):575. Epub 2022/10/31. doi: 10.1007/s00018-022-04607-5. PubMed PMID: 36309617; PubMed Central PMCID: PMC9617967.

45. Vinokurov AY, Soldatov VO, Seregina ES, Dolgikh AI, Tagunov PA, Dunaev AV, et al. HPRT1 Deficiency Induces Alteration of Mitochondrial Energy Metabolism in the Brain. Mol Neurobiol. 2023;60(6):3147–57. Epub 2023/02/22. doi: 10.1007/s12035-023-03266-2. PubMed PMID: 36802322; PubMed Central PMCID: PMC10122629.

46. Sekine M, Fujiwara M, Okamoto K, Ichida K, Nagata K, Hille R, et al. Significance and amplification methods of the purine salvage pathway in human brain cells. J Biol Chem. 2024;300(8):107524. Epub 2024/07/04. doi: 10.1016/j.jbc.2024.107524. PubMed PMID: 38960035; PubMed Central PMCID: PMC11342100.

47. Deng Z, Ou H, Ren F, Guan Y, Huan Y, Cai H, et al. LncRNA SNHG14 promotes OGD/R-induced neuron injury by inducing excessive mitophagy via miR-182-5p/BINP3 axis in HT22 mouse hippocampal neuronal cells. Biol Res. 2020;53(1):38. Epub 2020/09/12. doi: 10.1186/s40659-020-00304-4. PubMed PMID: 32912324; PubMed Central PMCID: PMC7488096.

48. Lee Y, Lee J, Jo D-G. A novel function of CD44 in the pathogenesis of Alzheimer’s disease. Alzheimer’s & Dementia. 2023;19(S21):e076285. doi: 10.1002/alz.076285.

49. Yu H, Lin L, Zhang Z, Zhang H, Hu H. Targeting NF-κB pathway for the therapy of diseases: mechanism and clinical study. Signal Transduct Target Ther. 2020;5(1):209. Epub 2020/09/23. doi: 10.1038/s41392-020-00312-6. PubMed PMID: 32958760; PubMed Central PMCID: PMC7506548.

50. Cheignon C, Tomas M, Bonnefont-Rousselot D, Faller P, Hureau C, Collin F. Oxidative stress and the amyloid beta peptide in Alzheimer’s disease. Redox Biology. 2018;14:450–64. doi: 10.1016/j.redox.2017.10.014.

51. You H, Tsutsui S, Hameed S, Kannanayakal TJ, Chen L, Xia P, et al. Aβ neurotoxicity depends on interactions between copper ions, prion protein, and N-methyl-D-aspartate receptors. Proc Natl Acad Sci U S A. 2012;109(5):1737–42. Epub 2012/02/07. doi: 10.1073/pnas.1110789109. PubMed PMID: 22307640; PubMed Central PMCID: PMC3277185.

52. Bulcke F, Santofimia-Castaño P, Gonzalez-Mateos A, Dringen R. Modulation of copper accumulation and copper-induced toxicity by antioxidants and copper chelators in cultured primary brain astrocytes. J Trace Elem Med Biol. 2015;32:168–76. Epub 2015/08/26. doi: 10.1016/j.jtemb.2015.07.001. PubMed PMID: 26302925.

53. Zeki AA, Yeganeh B, Kenyon NJ, Ghavami S. Editorial: New Insights into a Classical Pathway: Key Roles of the Mevalonate Cascade in Different Diseases (Part II). Curr Mol Pharmacol. 2017;10(2):74–6. Epub 2017/04/26. doi: 10.2174/187446721002170301204357. PubMed PMID: 28440195; PubMed Central PMCID: PMC6018051.

54. Varma VR, Büşra Lüleci H, Oommen AM, Varma S, Blackshear CT, Griswold ME, et al. Abnormal brain cholesterol homeostasis in Alzheimer’s disease—a targeted metabolomic and transcriptomic study. npj Aging and Mechanisms of Disease. 2021;7(1):11. doi: 10.1038/s41514-021-00064-9.

55. Pappolla MA, Refolo L, Sambamurti K, Zambon D, Duff K. Hypercholesterolemia and Alzheimer’s Disease: Unraveling the Connection and Assessing the Efficacy of Lipid-Lowering Therapies. J Alzheimers Dis. 2024;101(s1):S371–s93. Epub 2024/10/18. doi: 10.3233/jad-240388. PubMed PMID: 39422957.

56. Coscueta ER, Sousa AS, Reis CA, Pintado MM. Phenylethyl Isothiocyanate: A Bioactive Agent for Gastrointestinal Health. Molecules. 2022;27(3). Epub 2022/02/16. doi: 10.3390/molecules27030794. PubMed PMID: 35164058; PubMed Central PMCID: PMC8838155.

57. Jaafaru MS, Abd Karim NA, Enas ME, Rollin P, Mazzon E, Abdull Razis AF. Protective Effect of Glucosinolates Hydrolytic Products in Neurodegenerative Diseases (NDDs). Nutrients. 2018;10(5). Epub 2018/05/09. doi: 10.3390/nu10050580. PubMed PMID: 29738500; PubMed Central PMCID: PMC5986460.

58. Asif M, Kala C, Gilani S, Imam S, Taleuzzaman M, Naaz F, et al. Protective Effects Of Isothiocyanates Against Alzheimer’s Disease. Current Traditional Medicine. 2021;07. doi: 10.2174/2215083807666211109121345.

59. Ross IA. Neurodegenerative Diseases. In: Ross IA, editor. Plant-Based Therapeutics, Volume 2: The Brassicaceae Family. Cham: Springer Nature Switzerland; 2024. p. 261-314.

